# Targeting serine dehydratase supports amino acid homeostasis and skin repair

**DOI:** 10.1101/2025.10.23.683531

**Authors:** Emeline Joulia, Ethan L. Ashley, Patrick Tseng, Citlalic Arechiga, Yuqiong Liang, Aurélie Laguerre, Maureen L. Ruchhoeft, Joseph Rainaldi, Michal K. Handzlik, Regis J. Fallon, Hyun Young Kim, Prashant Mali, David A. Brenner, Tatiana Kisseleva, Ye Zheng, Marin L. Gantner, Christian M. Metallo

## Abstract

Serine and glycine are altered in patients with metabolic disorders, and this dysregulation can lead to diverse pathologies^1–6^. Modulation of serine levels via diet can influence relevant phenotypes in mouse models of metabolic syndrome^7,8^. Here we identify serine dehydratase (Sds), a gluconeogenic hepatic enzyme involved in serine and threonine catabolism, as a key regulator of systemic serine and sphingolipid metabolism. We show that SDS is expressed and active in human liver tissue. Furthermore, Sds abundance strongly correlates with hepatic serine. This enzyme is highly active in BKS-db/db mice, which show amino acid alterations reminiscent of type 2 diabetes. Hepatic Sds overexpression increases serine and threonine degradation and promotes the accumulation of toxic 1-deoxysphingolipids (doxSLs). Conversely, Sds deletion dramatically increases systemic serine, glycine, and threonine while altering canonical and non-canonical sphingolipids. Finally, Sds deletion in BKS-db/db mice reduces skin doxSLs and accelerates wound healing. Our results demonstrate that Sds constrains serine levels in circulation and suggest therapeutic approaches for targeting this enzyme to improve chronic disorders.

## Introduction

Diabetes is driven by both genetic and environmental factors that result in altered metabolic states caused by insulin resistance or deficiency, including aberrant carbohydrate, lipid, and amino acid metabolism. Heterogeneity in the pathogenesis of diabetes means patients experience a range of co-morbidities downstream of disease-associated metabolic dysregulation, encompassing renal failure, peripheral sensory neuropathy, impaired wound healing, and other pathologies^9^. Beyond dysregulated glucose, clinical metabolomics studies have demonstrated that serine, glycine, and threonine are reduced in subsets of diabetic patients, while branched-chain amino acids (BCAAs) are increased in the context of insulin resistance^10–12^. These changes may be linked to specific co-morbidities that can be targeted to improve health outcomes in patients via diet or other approaches. While circulating levels of serine and glycine are lower in several related metabolic diseases such as type 2 diabetes (T2D)^1,7^, metabolic dysfunction-associated steatotic liver disease (MASLD)^2–4^, and chronic kidney disease^5,6,13^, the key control points of amino acid homeostasis must be elucidated to facilitate therapeutic targeting. As genetic and physiological data are increasingly used to stratify subclasses of diabetic patients^14^, understanding the molecular drivers that dictate these metabotypes will become important.

Serine and glycine are highly exchanged non-essential amino acids (NEAAs) that have pleiotropic roles in metabolism that include sphingolipid biosynthesis^15–17^, nucleotide production^18,19^, methionine cycle^20–22^, redox metabolism^23–25^, and gluconeogenesis^7^. Serine and glycine homeostasis is maintained by the liver and kidney via one carbon metabolism. However, we recently observed that serine absorption is significantly reduced in mouse models of T2D, with a large fraction of serine converted to glucose in healthy animals^7^. The primary enzyme involved in this process is serine dehydratase (Sds), a hepatic enzyme which deaminates serine or threonine to yield pyruvate or alpha-ketobutyrate, respectively. The expression and activity of this enzyme is regulated in a manner consistent with its role in gluconeogenesis^26,27^. Nevertheless, little is known about its roles in coordinating amino acid and tissue homeostasis.

Numerous pathways may be impacted when serine levels are low. Sphingolipids are constitutively synthesized and catabolized in most tissues. Proper canonical sphingolipid production requires adequate availability of serine to sustain flux through the serine palmitoyltransferase (SPT) complex. When serine becomes limited, alanine may be used as a substrate for the reaction, yielding 1-deoxysphingolipids (doxSLs). DoxSLs cannot be degraded by the canonical phosphorylation pathway, and their accumulation can compromise cellular processes such as cell migration, vesicle trafficking, and anchorage-independent growth^28,29^. The nervous system and skin are highly enriched in sphingolipids, and their sustained biosynthesis and catabolism is critical for the function of these tissues. Our prior studies have demonstrated that serine availability reshapes the sphingolipidome and improves intraepidermal nerve fiber density in diabetic mice^7^. These data indicate that sphingolipid metabolism is highly sensitive to serine levels, and approaches that increase systemic serine could mitigate diabetic co-morbidities in peripheral tissues.

Here we describe the role of Sds in controlling serine and sphingolipid metabolism. We demonstrate that SDS is active in human liver tissue and correlates inversely with hepatic serine and threonine levels. Elevated Sds activity and serine/threonine clearance are also evident in diabetic mice, and Sds overexpression decreases serine, glycine, and threonine levels while promoting doxSLs synthesis. Conversely, Sds deletion drastically increases systemic serine, glycine, and threonine levels. However, sphingolipid biosynthesis is the predominant pathway altered by Sds deletion, significantly reducing the pool of doxSLs in liver and peripheral tissue. Finally, deletion of Sds in db/db mice results in elevated serine, glycine, and threonine along with decreased doxSLs in skin. This diminution in toxic sphingolipids in peripheral tissues is associated with improved wound healing. Collectively, these findings highlight the role of Sds in shaping serine and sphingolipid metabolism and establish this enzyme as a potential target to mitigate pathophysiology associated with low serine and/or glycine.

## Results

### Serine dehydratase is active in human livers

Based on our prior studies in mice showing rapid clearance of serine^7^, we performed a serine tolerance test (STT) in healthy subjects and tracked amino acid dynamics thereafter. Serine levels peaked after 60 minutes and achieved high concentrations that rapidly declined (**Fig. 1a**). Alanine and glycine were also transiently elevated in response to serine administration (**Fig. 1b**). This pronounced increase in alanine suggested SDS was active upstream of alanine aminotransferase (ALT) (**Fig. 1c**). Next, we directly assessed hepatic SDS activity in liver tissue from subjects with metabolic disease and compared these data to amino acid levels from the same tissue. While amino acids were not significantly different across disease states within this cohort (**Extended Data Fig. 1a**), threonine, but not serine, showed a significant, negative correlation with SDS activity (**Fig. 1d,e**). SDS activity was not distinctly increased in tissue from these non-fasted diabetic or MASLD subjects (**Extended Data Fig. 1b**), potentially due to the postprandial enzymatic regulation^26^. However, SDS was active in all patient tissues and exhibited high activity in a subset of subjects.

**Fig. 1.**
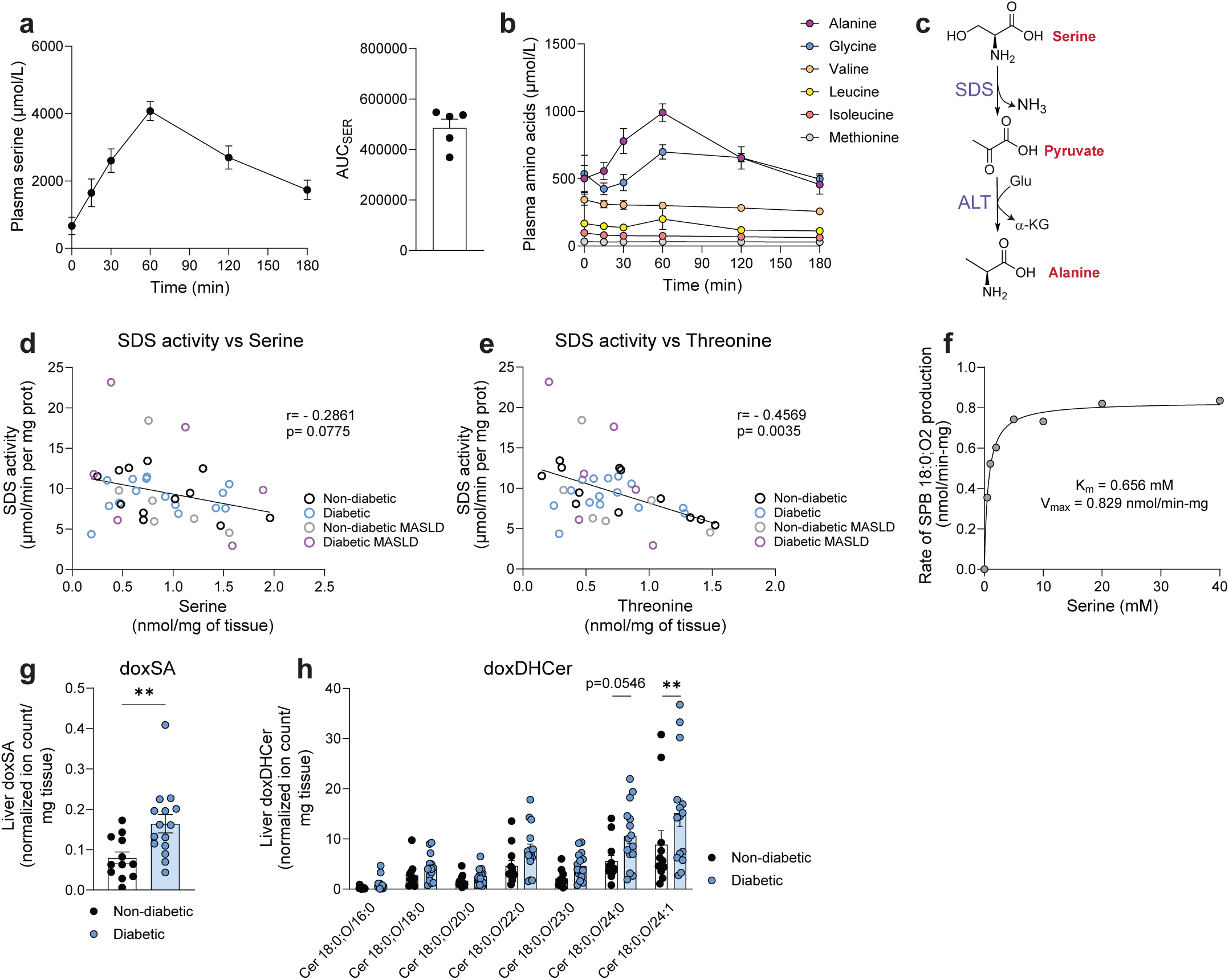
Human livers exhibit serine dehydratase activity. **a,b,** Plasma serine (**a**) and amino acid (**b**) concentration upon serine tolerance test performed in healthy subjects (n = 5). **c,** Alanine production from serine through serine dehydratase (SDS) and alanine aminotransferase (ALT). **d,e,** Correlation between SDS activity and serine (**d**) and threonine (**e**) concentrations in human livers (n = 39). **f,** Rate of sphinganine 18:0 production by microsomal SPT at varying concentrations of serine in HCT116 cells. **g,h,** doxSA 18:0 (**g**) and doxDHCer 18:0 (**h**) relative abundance in non-diabetic (n = 12) and diabetic livers (n = 15). Data are presented as mean ± standard error of mean (SEM). Pearson’s correlation analysis was performed to assess linear associations between variables, and statistical significance was determined using two-tailed p-values (**d,e**). Statistical analyses were performed using Mann-Whitney non-parametric t-test (**g**) and two-way ANOVA with Bonferroni’s multiple comparisons test (**h**).

Serine is an important substrate for sphingolipid biosynthesis through the serine palmitoyltransferase (SPT) complex, which can also metabolize alanine to generate doxSLs (**Extended Data Fig. 1c**). Prior studies have demonstrated that doxSLs correlate strongly with circulating serine and alanine^16^, highlighting an association between amino acid homeostasis and SPT flux. To better understand the biochemical link between serine levels and SPT flux, we measured the K_m_[Ser] for SPT in microsomes isolated from human cells. Consistent with reported values^17,30^, we measured the K_m_[Ser] of SPT to be over 0.6 mM (**Fig. 1f**), significantly higher than levels observed for circulating and hepatic serine. These data indicate that SPT is physiologically tuned to serine availability and indicate deoxydihydroceramides (doxDHCer) may be a longitudinal biomarker of this amino acid, analogous to HbA1C for glucose in diabetic subjects. In fact, we observed a significant increase in 1-deoxysphinganine (doxSA) and doxDHCer containing very long-chain fatty acids in diabetic subjects (**Fig. 1g,h**). These results are consistent with doxSL data in diabetic patients^1,31^. DoxSLs were generally high in subjects with MASLD but showed variability, potentially due to fat deposits (**Extended Data Fig. 1d,e**). Taken together, these data confirm that SDS is active in human liver and suggest that modulating its activity could influence doxSLs, a biomarker of aberrant serine and sphingolipid metabolism that is elevated in diabetic subjects.

### Serine dehydratase activity and expression are increased in diabetic mice

Next, to enable more direct metabolic analyses and genetic interventions in tissues, we examined this pathway in diabetic mice. To this end, we compared amino acid metabolism in leptin receptor-deficient mice, BKS-*Leprdb* (hereafter db/db mice), and BKS mice (WT) as control. Analysis of fasting amino acids revealed that systemic serine, glycine, methionine, and threonine were all significantly decreased in diabetic mice as compared to BKS controls (**Fig. 2a**). Conversely, branched-chain amino acids (BCAA) were increased (**Fig. 2a**), consistent with human data^12^. Intriguingly, threonine was the most decreased amino acid in this comparison and was associated with elevated α-ketobutyrate levels in diabetic mice (**Extended Data Fig. 2a,b**). These results highlight dysregulation of both one carbon metabolism and Sds in this diabetic mouse model. We then confirmed that serine clearance was elevated in diabetic mice administered a serine bolus (**Fig. 2b**). Serine catabolism was increased in diabetic mice that also exhibited increased pyruvate levels in the plasma upon STT, confirming increased Sds activity in db/db mice (**Fig. 2b,c**). In contrast to humans, little to no increase in alanine was observed after STT, highlighting a distinction between mouse and human physiology (**Extended Data Fig. 2c**). Moreover, by administering a bolus of threonine, we confirmed that threonine catabolism was also increased in db/db mice (**Extended Data Fig. 2d**). To validate this response was specific to serine and threonine, we performed an alanine tolerance test, observing no impact on alanine absorption or catabolism in db/db versus WT mice (**Extended Data Fig. 2e**). We then analyzed Sds expression and activity in control and db/db mouse livers and showed higher levels of both in diabetic mice, supporting its role in the elevated serine and threonine catabolism we measured in this mouse model (**Fig. 2d, Extended Data Fig. 2f**).

**Fig. 2.**
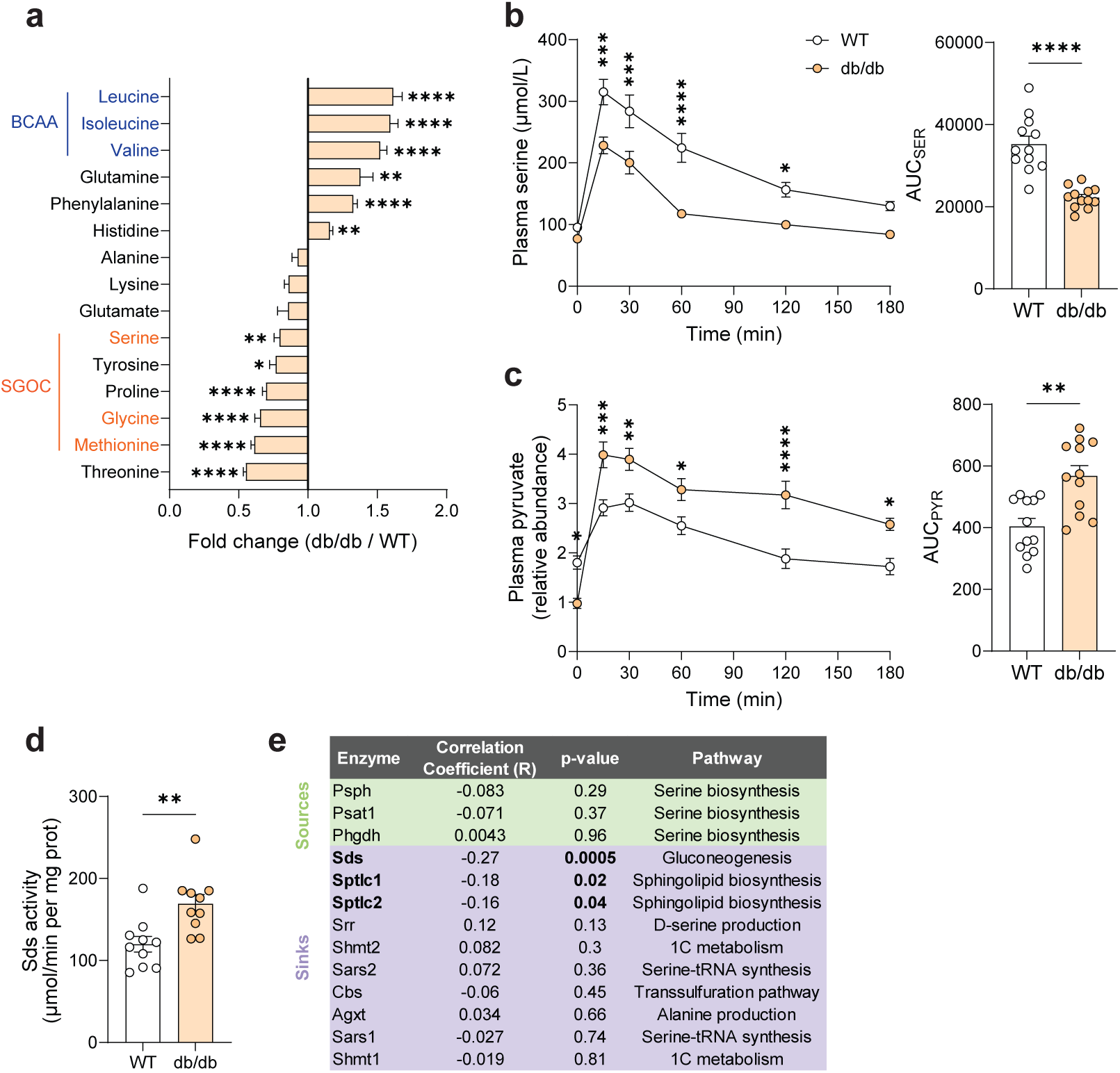
Diabetic mice exhibit increased serine dehydratase activity. **a,** Plasma amino acid profile in 10-14 week-old BKS-db/db mice (db/db) as compared to BKS-WT mice (WT) (n = 17 per group). **b,c,** Plasma serine concentration (**b**) and pyruvate relative abundance (**c**) upon serine tolerance test in WT and db/db mice (n = 12 per group). **d,** Sds activity in WT and db/db mouse livers (n = 10 per group). **e,** Correlation between serine metabolism-related enzyme expression and serine abundance in mouse liver from Xiao et al., 2025 (n = 163) - Psph: phosphoserine phosphatase, Psat1: phosphoserine aminotransferase 1, Phgdh: phosphoglycerate dehydrogenase, Sds: serine dehydratase, Sptlc1 and 2: serine palmitoyltransferase long chain base subunit 1 and 2, Srr: serine racemase, Shmt1 and 2: serine hydroxymethyltransferase 1 and 2, Sars1 and 2: seryl-tRNA synthetase 1 and 2, Cbs: cystathionine beta-synthase, Agxt: alanine-glyoxylate aminotransferase. Data are presented as mean ± standard error of mean (SEM). Statistical analyses were performed using two-way ANOVA with Bonferroni’s multiple comparisons test for multiple comparisons (**a**, **b** left, **c** left) and Mann-Whitney non-parametric t-test (**b** right, **c** right, **d**).

To assess what enzymes and pathways correlate most strongly with hepatic serine levels beyond db/db mice, we leveraged a unique database generated from liver metabolomics and proteomics measurements in diversity outbred (DO) mice^32^. Specifically, we investigated the relationship between enzymes involved in serine metabolism and abundance in mouse liver. Interestingly, Sds stood out as the most significant enzyme correlating with serine abundance (**Fig. 2e**). Notably, enzymes involved in glucose-driven synthesis showed no correlation with hepatic serine levels. Intriguingly, the only other enzymatic “source” or “sink” of serine that showed a significant correlation with serine abundance was sphingolipid biosynthesis. These data are consistent with the role of serine in sphingolipid production and the responsiveness of this pathway to dietary manipulation of serine^7,8^. Collectively, these findings demonstrate that Sds expression and activity are critically important for serine, glycine, and threonine metabolism in the db/db mice and other strains.

### Sds constrains circulating serine levels and influences sphingolipid metabolism

While early biochemical studies demonstrated the involvement of Sds in gluconeogenesis in liver^26,33^, the contribution of Sds to amino acid homeostasis has not been investigated in detail. To better understand the function of this enzyme, we first overexpressed SDS in the HCT116 cell line using lentivirus-based delivery and verified the presence of SDS by western blot (**Extended Data Fig. 3a**). SDS overexpression induced a significant decrease in serine, glycine, and threonine in these cells (**Extended Data Fig. 3b**), which actively converted [^13^C_3_]serine and [^13^C_2_]glycine to M+3 pyruvate (**Extended Data Fig. 3c**). No other changes in amino or organic acids were evident from SDS overexpression (not shown). On the other hand, several doxDHCer species were increased in these cells (**Extended Data Fig. 3d**), while canonical ceramides were not drastically altered after SDS overexpression (**Extended Data Fig. 3e**).

Next, we overexpressed Sds in C57Bl/6J mice using adeno-associated virus (AAV)-mediated delivery. We confirmed that increased Sds activity and expression in the liver was sustained 4 weeks after AAV injection (**Fig. 3a, Extended Data Fig. 3f**). Notably, Sds overexpression (Sds-OE) in mice induced a decrease in circulating serine, glycine, and threonine (**Fig. 3b**), and these mice showed reduced AUC_SER_ compared to NTC mice upon administration of a STT (**Fig. 3c**). We also observed a decrease in serine, glycine, and threonine (**Fig. 3d**) along with increased pyruvate and α-ketobutyrate in hepatic tissue (**Extended Data Fig. 3g,h**). Finally, we examined the impact of Sds overexpression on hepatic sphingolipid metabolism. 4 weeks after AAV injection, Sds overexpression induced a strong increase in doxDHCer in mouse liver (**Fig. 3e**) but caused no change in hepatic ceramides (**Extended Data Fig. 3i**). These data indicate that hepatic Sds overexpression alters systemic serine, glycine, threonine, and doxSL levels in mice.

**Fig. 3.**
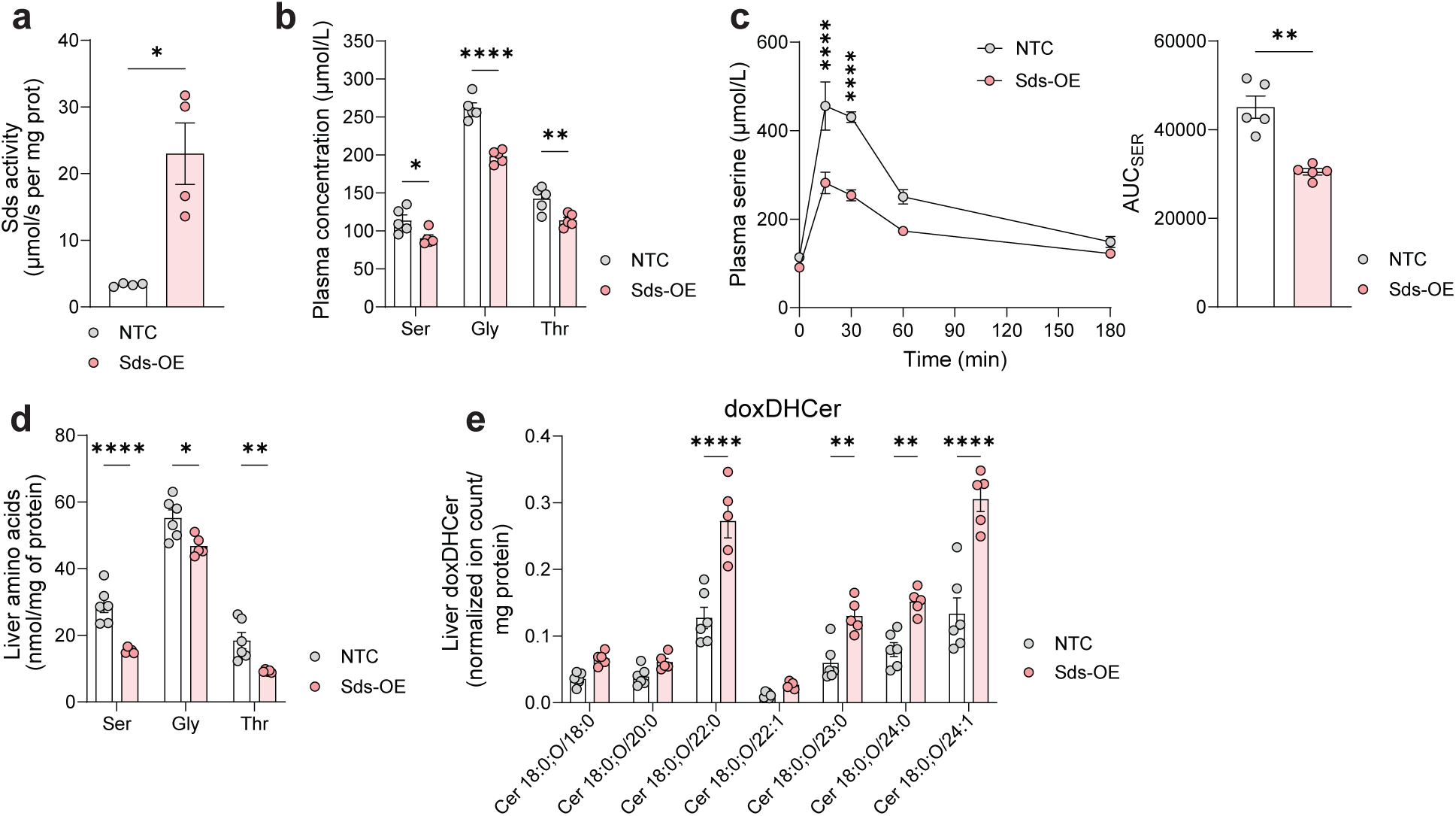
Sds overexpression promotes serine catabolism and increases non-canonical sphingolipids. **a,** Sds activity in mouse liver, 4 weeks post-AAV injection (n = 4 per group). **b,c,** Plasma amino acid profile (**b**) and serine tolerance test (**c**) 10 days post-AAV injection in NTC and Sds-OE mice (n = 5 per group). **d,e,** Hepatic amino acid (**d**) and deoxydihydroceramides 18:0 (**e**) profile in NTC (n = 6) and Sds-OE mouse livers (n = 5), 4 weeks post-AAV injection. Data are presented as mean ± standard error of mean (SEM). Statistical analyses were performed using Mann-Whitney non-parametric t-test (**a, c** right) and two-way ANOVA with Bonferroni’s multiple comparisons test (**b**, **c** left, **d** and **e**).

### Sds deletion metabolically supports amino acid and sphingolipid metabolism

Next, we obtained a Sds knockout (KO) mouse model in C57BL/6 (B6) background to better understand the contribution of this enzyme to amino acid physiology. We first confirmed that Sds was undetectable by western blot and enzyme assay in the liver, the only tissue showing appreciable expression or activity in WT mice (**Extended Data** Fig. 4a,b). We then quantified amino acids in circulation and observed that serine, glycine and threonine were increased by 5-, 2.9- and, 5.8-fold, respectively, in fasted Sds-KO mice (**Fig. 4a,b**). Upon administration of a STT, we noted that Sds deletion drastically increased serine and glycine levels, which were sustained at very high levels for over 3 hours after the STT was initiated (**Fig. 4c, Extended Data Fig. 4c**). Given the striking and prolonged increase in amino acids observed in Sds-KO mice after consuming serine, these data suggest Sds is the primary means through which mice constrain supraphysiological serine, glycine, and threonine.

**Fig. 4.**
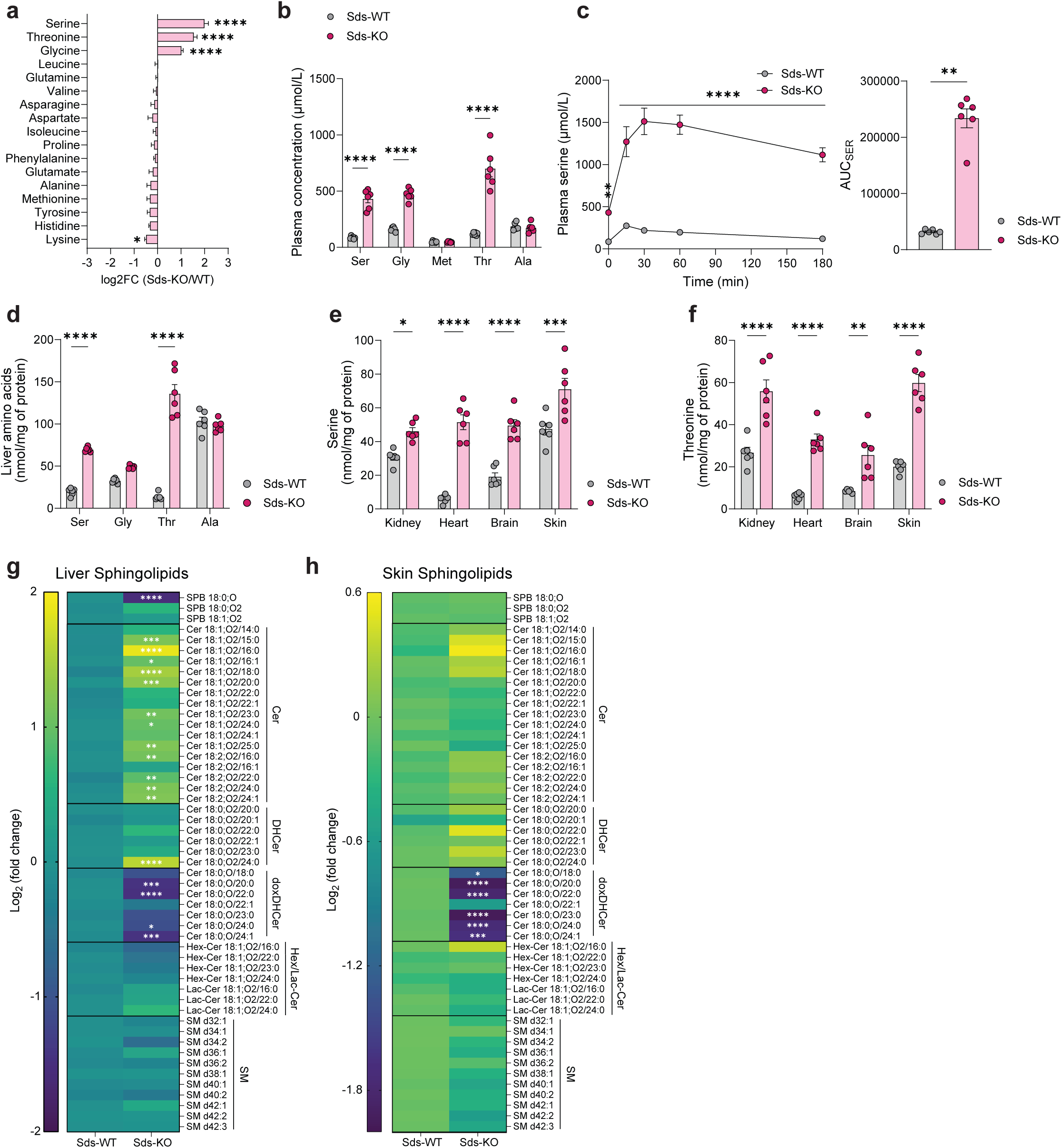
Serine dehydratase deletion mainly affects serine, threonine and sphingolipid metabolism. **a,** Circulating amino acid profile in Sds-WT (n = 18) vs Sds-KO mice (n = 14). **b,** Plasma serine, glycine, methionine, threonine and alanine concentration in Sds-WT and Sds-KO mice (n = 6 per group). **c,** Serine tolerance test performed on 8-week-old mice (n = 6 per group). **d,** Amino acid concentrations in Sds-WT versus Sds-KO mouse livers (n = 6 per group). **e,f,** Serine (**e**) and threonine (**f**) concentration in kidney, heart, brain and skin from 10-12 week-old Sds-WT and KO mice (n = 6 per group). **g,h,** Liver (**g**) and skin (**h**) sphingolipids in 10–12-week-old mice (n = 6 per group) - Cer: Ceramides, DHCer: Dihydroceramides, doxDHCer: deoxydihydroceramides, Hex/Lac-Cer: Hexosyl/Lactosyl-Ceramides, SM: Sphingomyelin. Data are presented as mean ± standard error of mean (SEM). Statistical analyses were performed using two-way ANOVA with Bonferroni’s multiple comparisons test (**a**, **b**, **c** left, **d-h**) and Mann-Whitney non-parametric t-test (**c** right).

Consistent with amino acid changes in blood, increased serine and threonine were also observed in Sds-KO mouse liver (**Fig. 4d**). Furthermore, Sds deletion had a systemic impact on amino acid metabolism, as serine and threonine were elevated in a range of tissues, including kidney, heart, brain and skin (**Fig. 4e,f**). Since serine contributes to numerous metabolic pathways, we examined whether any of these downstream pathways were altered significantly in the liver (**Extended Data Fig. 4d**). Despite the role of Sds in gluconeogenesis, no changes were evident in fasting or fed blood glucose (**Extended Data Fig. 4e**), indicating serine and threonine contribute minimally to glucose pools during normal physiology. Glucose tolerance was also unaffected by Sds deletion (**Extended Data Fig. 4f**). Similarly, we measured metabolites in the glutathione and purine synthesis pathways and observed no significant changes (**Extended Data Fig. 4g-i**). On the other hand, modest increases in hepatic methionine and cysteine metabolism were detected, suggesting increased serine may support the transsulfuration pathway (**Extended Data Fig. 4j**). L-serine is also converted to D-serine by serine racemase, primarily in the brain and kidney^34,35^ (**Extended Data Fig. 4d**). Plasma D-serine levels were dramatically increased in Sds-KO mice (**Extended Data Fig. 4k**), suggesting elevated kidney disposal in the absence of Sds. Finally, we analyzed sphingolipids in Sds-WT and KO mice. Sds deletion strongly reduced doxSLs in both hepatic and peripheral (skin) tissue (**Fig. 4g,h**). On the other hand, canonical ceramides in the liver were elevated in Sds-KO mice, with abundant very long-chain ceramides (24:0 and 24:1) being the most impacted (**Extended Data Fig. 4l)**. No major differences were noted in sphingomyelins (SM) or simple glycosphingolipids (Hex/Lac-Cer) in both liver and skin (**Fig. 4g,h**). Altogether, these results demonstrate that Sds sets the upper limit for circulating serine (and threonine) in mice, which primarily impacts downstream sphingolipid metabolism in hepatic and peripheral tissues. Targeting this pathway could therefore support amino acid and sphingolipid homeostasis systemically.

### Sds deletion improves wound healing in diabetic mice

Diabetes is associated with co-morbidities such as peripheral neuropathy and impaired wound healing. We previously observed that modulating dietary serine availability and doxDHCer synthesis influences nerve fiber innervation in the epidermis^7^, suggesting that serine is important in supporting peripheral tissue homeostasis. Indeed, doxSLs are increased in diabetic mice and humans^1,7,16,36^, and these metabolites can influence anchorage-independent growth, cell migration, and other processes important for tissue regeneration^28,29^. To investigate whether targeting Sds could have a beneficial impact on wound healing in diabetic mice, we generated db/db-Sds-KO mice by crossing B6-Sds-KO mice with db/db-BKS heterozygotes. We first analyzed whether Sds deletion affected body weight and blood glucose levels during growth from 3- to 7-week-old. Sds deletion did not impact either parameter, demonstrating that Sds deficiency does not affect diabetes progression in this mouse model (**Extended Data Fig. 5a,b**). We also performed a glucose tolerance test on both male and female db/db-Sds-WT and KO mice, observing no impact on glucose clearance (**Extended Data Fig. 5c**). Next, we analyzed the amino acid profile in the plasma and liver. db/db-Sds-KO mice exhibited increased serine, glycine, and threonine both in circulation and liver (**Fig. 5a,b**). Importantly, db/db-Sds-KO mice given a STT had significantly elevated serine and glycine levels in circulation that were sustained for several hours after serine administration (**Fig. 5c, Extended Data Fig. 5d**).

**Fig. 5.**
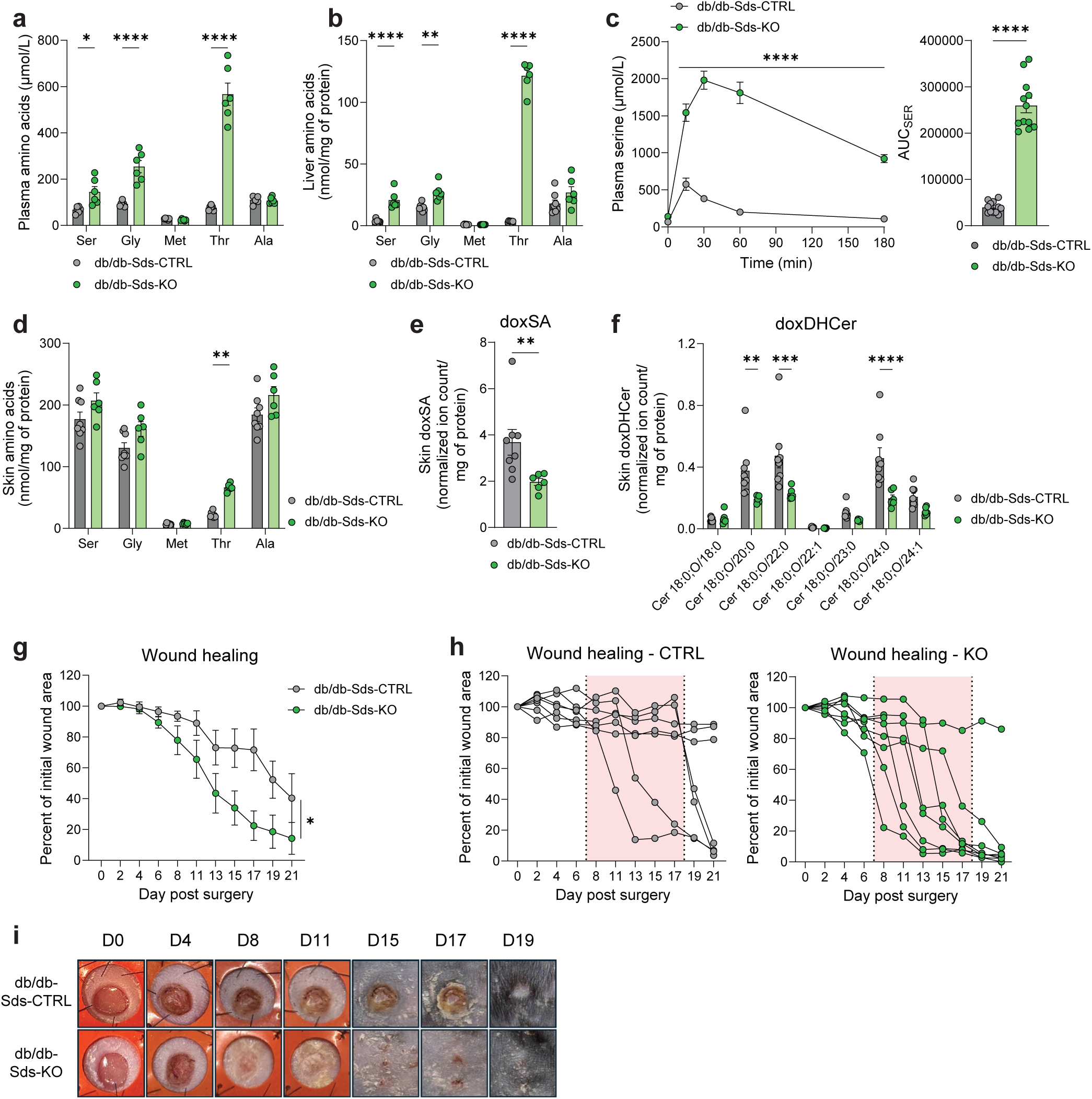
Serine dehydratase deletion restores serine/glycine and sphingolipid metabolism and improves wound healing in diabetic mice. **a,b,** Plasma (**a**) and liver (**b**) amino acid concentration in db/db-Sds-WT (n = 8) and db/db-Sds-KO mice (n = 6). **c,** Serine tolerance test performed on 10-14-week-old female and male mice (n = 12 per group). **d,** Skin amino acid concentration in db/db-Sds-WT (n = 8) and db/db-Sds-KO mice (n = 6). **e,f,** Deoxysphinganine 18:0 (**e**) and deoxydihydroceramide relative abundance (**f**) in db/db-Sds-WT (n = 8) and db/db-Sds-KO (n = 6) mouse skin. **g,** Wound healing measurement in db/db-Sds-CTRL (n = 7) versus db/db-Sds-KO (n = 8) until day 21 post-surgery. **h,** Individual representation of wound closure over 21 days in db/db-Sds-CTRL (n = 7) and db/db-Sds-KO mice (n = 8). **i,** Representative pictures upon healing process from day 0 to day 19 post surgery. Data are presented as mean ± standard error of mean (SEM). Statistical analyses were performed using two-way ANOVA with Bonferroni’s multiple comparisons test (**a**, **b**, **c** left, **d, f** and, **g**) and Mann-Whitney non-parametric t-test (**c** right, **e**).

We next measured amino acids in peripheral skin tissue. Here only threonine levels were significantly elevated, and we observed no impact on serine and glycine levels, likely due to their higher abundance in skin versus liver (**Fig 5b,d**). On the other hand, doxSA and doxDHCer were reduced in the skin of db/db-Sds-KO mice by approximately 50% (**Fig. 5e,f**), similar to findings in patients^37^ and db/db mice fed a serine-enriched diet^7,8^. No other sphingolipids were significantly altered in db/db-Sds-KO mouse skin (**Extended Data Fig. 5e**). These results suggest that elevated serine caused by Sds deficiency improves the regenerative microenvironment by reducing doxSLs in skin. We therefore hypothesized that db/db-Sds-KO mice might exhibit enhanced wound healing via improved amino acid and sphingolipid homeostasis. In mice, since wound healing relies on skin contraction rather than cell proliferation, we used the splint model^38^ to assess wound closure kinetics. Intriguingly, we observed that Sds deletion improved wound healing in diabetic mice (**Fig. 5g**), with 7/8 db/db-Sds-KO mice recovering between day 7 and day 18 post surgery against only 2/7 healing completely in the control group (**Fig. 5h,i**). These data indicate that targeting Sds could enhance amino acid and sphingolipid metabolism that becomes compromised in patients with diabetes or other chronic diseases. By limiting the catabolism of non-essential amino acids critical for cell growth and membrane biosynthesis, metabolic deficiencies induced by diabetes could be mitigated to improve wound healing or other co-morbidities.

## Discussion

Here we describe Sds as a potential target for regulating systemic amino acid and sphingolipid metabolism. Deleting Sds in both non-diabetic and diabetic mice reduces serine and threonine catabolism, increases serine, glycine, and threonine availability, and modulates the sphingolipid profile in both liver and peripheral tissues. A major consequence of these changes is a reduction of doxSLs, which are associated with reduced cell migration, impaired cell growth, and other defects^28,29^. By generating db/db-Sds-KO mice, we linked these metabolic changes to improved wound healing in diabetic skin.

Although our Sds deletion models are full-body KO, our results along with human and mouse transcriptomic data indicate the enzyme is mainly expressed in hepatocytes^39–41^. Remarkably, eliminating this hepatic activity has little effect on glucose but dramatic effects on systemic serine, glycine, and threonine levels. These findings highlight a unique role for Sds in constraining the upper limits of these amino acids in circulation. Evidently, Sds is the primary route for supraphysiological serine and threonine disposal, as the KO animal seemingly relies on kidney efflux to reduce amino acid levels after a STT. Sds therefore plays an important role in regulating serine and sphingolipid fluxes in diverse tissues. Indeed, our SPT K_m_[Ser] data further demonstrate the responsiveness of this enzyme complex to physiological serine levels as well as the strength of doxDHCer as a longitudinal serine biomarker. Intriguingly, these data also suggest that ectopic Sds expression in the liver could be beneficial in instances where SPT flux is aberrantly high^42,43^.

Our data also demonstrate a major role for Sds in threonine catabolism, as Sds-KO mice appear to rely exclusively on this enzyme for disposal. In some clinical metabolomics studies of patients with metabolic syndrome^2^, threonine is reduced along with serine and glycine. This pathway was also identified in a recent study of caloric restriction-induced longevity in mice and non-human primates^44^. Altered SDS expression/activity could be responsible for these metabotypes, which are associated with both metabolic syndrome and aging. Unlike serine and glycine, threonine is an essential amino acid, and the impacts of its accumulation on mouse physiology remain unclear. Even though we did not observe any obvious phenotype in Sds-KO mice, given the role of threonine in mucus production^45^, muscle mass maintenance^46^ and fat deposition^47^, the consequences of threonine accumulation in Sds deficient mice warrant further investigation.

We have much to learn about Sds regulation. While there is clear regulation of this gluconeogenic enzyme by hormones like insulin and glucagon^26^, our data in liver tissues from subjects that are non-diabetic, diabetic, and/or have MASLD suggests more complex mechanisms may be at play. In each group, we identified subjects with higher SDS activity correlating with reduced serine and threonine concentration in the liver. While glucocorticoids have been suggested to drive Sds expression in rats^27^, more extensive clinical studies are required to understand this regulation in humans. Nevertheless, the enzyme is active in humans and likely influences circulating amino acid and sphingolipid levels with physiological consequences in some patients.

Diabetes co-morbidities are often considered to be a consequence of hyperglycemia, which can negatively impact diverse tissue functions^48,49^. Our data show that Sds improves wound healing in diabetic mice independently of any effect on glycemia, further highlighting a critical role of serine/glycine homeostasis in diabetes comorbidities. Our work is consistent with previous studies showing the benefit of dietary serine supplementation on diabetic peripheral neuropathy without affecting hyperglycemia^7,8^. Hence, regulating amino acid homeostasis should be further explored to address diabetes-associated co-morbidities in a glucose-independent manner.

From a therapeutic perspective, strategies to augment systemic serine (and/or glycine) have been explored in clinical trials and/or pre-clinical models encompassing numerous diseases, including cardiovascular disorders^50–53^, Alzheimer’s disease^54^, diabetic peripheral neuropathy^7,31^, macular telangiectasia (MacTel)^16^, pancreatitis^55^, colitis^56^, and wound healing/tissue regeneration^29,57–59^. Our serine tolerance studies in mice and humans demonstrate challenges in sustaining high levels of circulating serine and glycine via dietary supplements, since SDS potently reduces levels back to baseline. Targeting SDS with or without concurrent amino acid supplementation may therefore emerge as an effective means of mitigating pathophysiology associated with diabetes, aging, or genetic disorders.

## Methods

### Mouse experiments

Experimental protocols were approved and conducted in accordance with the Institutional Animal Care and Use Committee (IACUC) of the Salk Institute for Biological Studies. Seven-week-old C57BL/6J (#000664), C57BLKS/J (#000662) and BKS.Cg-Dock7m +/+ Leprdb/J (BKS-*db/db*) mice (#000642) were purchased from Jackson. C57BL/6-*Sds^tm1.2^ ^mrl^* (Sds) heterozygotes mice were purchased from Taconic (#10998) and Sds-WT and Sds-KO were obtained after several crossings. db/db-Sds-WT and KO mice were generated after crossing BKS-*db/db* heterozygotes mice with Sds-KO mice. Mice were housed at room temperature (21°C) in the same room with a 12-hour light-dark cycle with water access provided *ad libitum*. Mice were fasted overnight (∼15hrs) prior to amino acid tolerance test or tissue collection. Mice were anesthetized with isoflurane and tissues were rapidly collected using Wollenberger clamps pre-cooled to the temperature of liquid nitrogen and stored at -80°C until analysis. Blood was collected in EDTA-coated tubes (Sarstedt Inc.) and centrifuged at 2000g at 4°C for 5 minutes to collect the plasma. Samples were transferred to a new Eppendorf tube and stored at -80°C until analysis.

### Donor samples

De-identified donor livers (IRB 171883XX; certified by the HRPP Director and IRB Chair to be “no human subjects” under the Code of Federal Regulations, Title 45, part 46 and UC San Diego Standard Operating Policies and Procedures) were obtained from the Lifesharing OPO, which provided informed consent, laboratory data (liver biopsy, biochemistry, ALT, AST, cell counts, serology, etc.), and patient’s history (age, gender, BMI, comorbidities, and cause of death). Livers were graded by a pathologist in a double-blind manner and identified as NORMAL and MASL(≥5% steatosis). Donor diabetes status was determined from their medical history. Snap-frozen liver specimens were used for analysis as described below.

### Amino acid tolerance tests (Serine TT, Threonine TT, Alanine TT)

#### Mice

Age-matched eight to twelve-week-old mice were fasted overnight with water access provided *ad libitum*. Prior to any tolerance test, mice were weighted, and serine, threonine or alanine was administered via oral gavage at a dose of 400mg/kg of body weight with tail tip blood samples collected into EDTA-coated tubes (Sarstedt Inc.) at baseline and 15, 30, 60, 120 and 180 minutes after amino acid administration. Tubes were centrifuged at 2000g at 4°C for 5 minutes and plasma samples were collected and stored at -80°C until analysis.

#### Humans

Human subjects read and signed informed consent forms approved by our local Institutional Review Boards. This research is in compliance with the Declaration of Helsinki and HIPAA privacy regulations. Volunteers were fasted overnight (∼10hrs) before serine tolerance test. Body weights were collected, and serine was dissolved in drinking water at 400mg/kg of body weight. Finger pricks were performed at baseline and 15, 30, 60, 120 and 180 minutes post serine ingestion. Blood samples were collected with capillaries and plasma was isolated and stored at -80°C until analysis.

### Glucose tolerance test (GTT)

Age-matched eight to twelve-week-old mice were fasted overnight (∼15hrs) with water provided *ad libitum*. In the morning, mice were weighed and fasting blood glucose determined from the tail bleed using Contour Next glucometer (Bayer). Next, glucose was administered by oral gavage at a dose of 2 g/kg of body weight, and blood glucose determined at 15, 30, 60 and 180 minutes post glucose administration.

### Wound healing assay

Wound healing assay was performed as previously described^38^. Briefly, mice were anesthetized, and the dorsal surface was shaved and applied with a depilatory agent to remove any remaining hair. A donut-shaped splint (0.5 mm-thick silicone splint, Grace Bio-Laboratories, Bend, OR) was placed on the mouse dorsal surface and an immediate-bonding adhesive (Krazy GlueÒ; Elmer’s Inc., Columbus, OH) was used to fix the splint to the skin followed by interrupted 6–0 nylon sutures (Ethicon, Inc., Somerville, NJ) to ensure position. A sterile 4-mm punch biopsy tool was used to outline a pattern for the wounds on the dorsum of the db/db mice, the tool was placed so that the wound was center within the splint. Full-thickness wounds extending through the panniculus carnosus were made using an Iris scissor. The animals were single housed in the animal facility. Digital photographs were taken on the day of surgery and every other day thereafter. Wound closure was measured using QuPath software (version 0-5-1).

### Adeno-associated virus (AAV) and lentivirus plasmid production

To generate lentiviral and AAV overexpression constructs, gene fragments encoding mouse Sds (mSds) and human SDS (hSDS) were synthesized as gBlocks (IDT) and cloned into the lentiviral vector pEPIP (Addgene #172110) by Gibson Assembly using the EcoRI multiple cloning site. The same fragments were also Gibson assembled downstream of the CMV promoter into the AAV plasmid backbone pZac2.1 (University of Pennsylvania Vector Core). For control constructs, eGFP was cloned into the EcoRI site of pEPIP, and mCherry was cloned into pZac2.1, using the same approach. AAV and lentivirus plasmid sequences were listed in **Supplementary Table 1**.

### AAV production and mouse injection

Adeno-associated viruses serotype 9 (AAV9) were generated by the Gene Transfer, Targeting and Therapeutics (GT3) Viral Core at the Salk Institute for Biological Studies (La Jolla, CA). 8-week-old C57BL/6J mice (Jackson, #000664) were injected with 0.7E+12 genome copy/mouse intravenously, by retro-orbital injection.

### Lentivirus production and SDS OE stable cell line generation

Cell lines overexpressing human SDS or a control EGFP sequence were produced by lentiviral transduction. Lentiviral particles were produced according to the Addgene pLKO vector protocol. In brief, lentiviral particles were packaged in HEK293T cells that were transfected with a pLKO transfer plasmid and the packaging plasmids psPAX2 (Addgene Plasmid #12260) and pMD2.G (Addgene Plasmid #12259) using FuGene 6 (FuGENE cat. no. F6-1000). 3 mL of DMEM supplemented with 10% FBS (Thermo Fisher Scientific) and 1% PenStrep (Thermo Fisher Scientific) was added to the cells at 24 and 48 hours post transfection and collected at 48 and 72 hours post transfection. The lentiviral suspension was filtered through a 0.45 μm filter to remove cellular debris and supplemented with polybrene to a concentration of 6 μg/mL.

HCT116 cells (∼250,000) were transduced with 500 μL of viral suspension. After 6 hours, they were supplemented with 2 mL of virus-free DMEM. The following day, DMEM containing 3 μg/mL puromycin (GoldBio) was added to select for cells that were successfully transduced.

### Serine dehydratase activity assay

The Sds activity assay was performed as described previously^7^. 25-30 mg of frozen liver tissue was extracted in ice cold buffer containing 50 mM KH2PO_4_, 1 mM Na_2_ EDTA, and 1 mM DTT, pH 8.0 with a glass homogenizer. Enzyme activity was determined using a coupled-enzyme reaction with 8.25 units of lactate dehydrogenase (Sigma 10127230001) per reaction in the presence of 0.25 mM NADH, 0.17 mM pyridoxal phosphate, and 1 mM DTT. The reaction was initiated by addition of serine to 200 mM and the absorbance of NADH was measured at 340 nm for up to 60 minutes. Protein quantification was performed by BCA protein assay and enzyme activity was expressed in units of specific rate (nmol substrate/minute/mg protein).

### Stable isotope tracing

NTC and SDS-OE HCT116 cells (∼300,000) were cultured in tracing media that was reconstituted from glucose, glutamine, and amino acid free DMEM (Sigma Aldrich cat. No. D5030) supplemented with ^13^C labeled serine and glycine (CLM-1574-H and CLM-1017-PK), unlabeled constituents, and 10% FBS. Serine and glycine were each supplemented to a concentration of 400 μM. Cells were washed once with PBS before addition of the tracing media. After 6 hours, tracing media was removed, and cells were washed once with 0.9% saline prior to metabolite extraction and quantification as described below.

### Metabolite extraction

#### Plasma

Plasma polar metabolites were extracted from 3 μL of plasma spiked with 300 pmol of ^13^C^15^N labeled amino acid standards (Cat# MSK-CAA-1, Cambridge Isotope Laboratories) and 300 pmol of DL-Norvaline (Cat#N7502, Sigma-Aldrich).

#### Cells

Cells (∼600,000) were spiked with the following internal standards: 20 pmol sphinganine-d7 (Avanti Polar Lipids, Cat# 860658), deoxysphinganine-d3 (Avanti Polar Lipids, Cat# 860474), 100 pmol d18:0-d7/13:0 dihydroceramide (Avanti Polar Lipids, Cat# 330726), 200 pmol d18:1-d7/15:0 ceramide (Avanti Polar Lipids, Cat# 860681), 100 pmol d18:1-d7/15:0 glucosylceramide (Avanti Polar Lipids, Cat# 330729), 100 pmol d18:1-d7/15:0 lactosylceramide (Avanti Polar Lipids, Cat# 330727), 200 pmol sphingosine d7(Avanti Polar Lipids, Cat# 860657), and 200 pmol d18:1/18:1-d9 sphingomyelin (Avanti Polar Lipids, Cat# 791649), and 500 pmol of ^13^C,^15^N-labelled amino-acid mixture (Cat# MSK-CAA-1, Cambridge Isotope Laboratories). Cells were scraped with 500 μL methanol and 500 μL water containing an additional 2.5 μg unlabeled DL-norvaline (Cat#N7502, Sigma-Aldrich) internal standard. 100 μL of homogenate was taken to determine protein content. 1 mL of chloroform was added to the remaining polar phase and then vortexed and centrifuged at 4 °C at 21,000g for 5 min each. The organic phase was collected and 2 μL of formic acid was added to the polar phase and then re-extracted with 1 mL of chloroform. The combined organic phases were dried under nitrogen and 300 μL of the polar phase was dried under vacuum.

#### Tissues

Tissue extraction was performed as described before^7^. Briefly, frozen tissues (15-20mg) were spiked with the following internal standards: 20 pmol sphinganine-d7 (Avanti Polar Lipids, Cat# 860658), deoxysphinganine-d3 (Avanti Polar Lipids, Cat# 860474), 100 pmol d18:0-d7/13:0 dihydroceramide (Avanti Polar Lipids, Cat# 330726), 200 pmol d18:1-d7/15:0 ceramide (Avanti Polar Lipids, Cat# 860681), 100 pmol d18:1-d7/15:0 glucosylceramide (Avanti Polar Lipids, Cat# 330729), 100 pmol d18:1-d7/15:0 lactosylceramide (Avanti Polar Lipids, Cat# 330727), 200 pmol sphingosine-d7 (Avanti Polar Lipids, Cat# 860657), and 200 pmol d18:1/18:1-d9 sphingomyelin (Avanti Polar Lipids, Cat# 791649), 2nmol of ^13^C^15^N labeled amino acid standards (Cat# MSK-CAA-1, Cambridge Isotope Laboratories) and 2nmol of DL-Norvaline (Cat#N7502, Sigma-Aldrich). For purine metabolites analysis, samples were spiked with 20 nmol of Piperazine-N,N′- bis(2-ethanesulfonic acid) (PIPES). Tissue was homogenized for 5 minutes using Precellys beads (#P000926-LYSK0-A.0) with 500 µL of -20°C methanol and 500 µL of HPLC grade H_2_O. Homogenate aliquot of 100 µL was taken to determine protein content using the BCA protein assay (Thermo Fisher Scientific). The remaining homogenate was transferred to a new Eppendorf tube and 1 mL chloroform was added. Samples were vortexed for 5 minutes and centrifuged for 5 minutes at 4°C at 21,000g. The organic phase was collected and 2 µL of formic acid was added to the remaining polar phase which was re-extracted with 1 mL of chloroform. Combined organic phases were dried under nitrogen. The polar phase was collected and dried under vacuum.

### Metabolite quantification

#### L-Amino acids and organic acids

Quantification of polar metabolites was determined as previously described^7^. Briefly, quantification was made after derivatization with 2% (w/v) methoxyamine hydrochloride (Thermo Scientific) in pyridine (37 °C for 60 min) and with N-tert-butyldimethylsilyl-N-methyltrifluoroacetamide (MTBSTFA) with 1% tert-butyldimethylchlorosilane (tBDMS) (Regis Technologies) (37 °C for 30 min). Polar derivatives were analyzed by GC–MS using a DB-35MS column (30 m × 0.25 mm i.d. × 0.25 μm, Agilent J&W Scientific) installed in an Agilent 7890 B gas chromatograph (GC) interfaced with an Agilent 5977 A mass spectrometer (MS) as previously described^7^. Natural isotope abundance was corrected using in-house script.

#### D-serine

D-serine quantification was performed using an Astec Chirobiotic T column (150 x 4.6 mm, 5 µm particles size, Supelco) installed on a triple quadrupole liquid chromatography mass spectrometry platform (6460, Agilent). Mobile phase was isocratic with 20% mobile phase A containing 100% HPLC-grade water with 0.1% formic acid, and 80% mobile phase B containing 100% methanol with a flow rate of 0.3 mL/min. Data were analyzed with MassHunter Workstation Software (Version B.08.00).

#### Purine metabolites

Purine metabolites were quantified using HILIC-HRMS with a Vanquish Flex UHPLC on an InfinityLab Poroshell 120 HILIC-z column (2.1 x 100mm, 2.7 µm, Agilent). Mobile phases composition, flow rate and gradient are detailed elsewhere^53^. Data were acquired via a Q-Exactive orbitrap mass spectrometer, with HESI conditions and MS parameters as previously described^60^. Concentrations were calculated using TraceFinder and a 10-point standard curve. Data were analyzed with El MAVEN Software (Version 0.11.0).

#### Sphingolipids

Sphingolipids were separated using a C8 column (Spectra 3 μm C8SR 150 x 3 mm inner diameter, Peeke Scientific) installed on a triple quadrupole liquid chromatography mass spectrometry platform (6460, Agilent). Dried extracts were resuspended in 80 μL buffer B and 60 μL was transferred to inserts in GC vials. 5 μL of the sample was injected. Mobile phases composition, flow rate and gradient are detailed elsewhere^7^. Sphingolipid species were analyzed by multiple reaction monitoring (MRM) of the transition from precursor to product ions at associated optimized collision energies and fragmentor voltages as previously described^61^. Relative abundance of sphingolipids was calculated by normalizing to internal standard specific to sphingolipid class and to protein levels. Data were analyzed with MassHunter Workstation Software (Version B.08.00).

### Sulfur metabolites extraction and quantification

Tissue extraction and quantification were performed as previously described with some modifications^62^. Briefly, frozen tissues (∼20mg) were spiked with 5nM of Piperazine-N,N′-bis(2-ethanesulfonic acid) (PIPES). Tissue was homogenized for 5 minutes using Precellys beads (#P000926-LYSK0-A.0) in 1 mL of freshly made ice cold extraction solution (80% MeOH with 20% 10 mM ammonium formate and 25 mM N-Ethylmaleimide (NEM)). Homogenate aliquot of 50 µL was taken to determine protein content using the BCA protein assay (Thermo Fisher Scientific). Samples were vortexed briefly and the remaining homogenate was transferred to a new Eppendorf tube. After 30 minutes of incubation on ice, samples were vortexed for 5 minutes and centrifuged for 5 minutes at 4°C at 21,000g. Supernatants were collected and dried under vacuum. Dried samples were resuspended in 100 µL of 80% MeOH and transferred to a vial for detection and quantification by liquid chromatography tandem mass spectrometry (LC-MS/MS). Metabolites quantification was performed using an isotope dilution method with both external standard curves and internal standards following the method previously described (Kambhampati et al., 2019), with modifications. Briefly, 3 µL of sample was injected onto an Infinity Lab Poroshell 120 Z-HILIC column (2.7 um, 100 x 2.1 mm; Agilent technologies, Santa Clara, CA, USA) installed on a liquid chromatography system (1260 Infinity II, Agilent) coupled to a triple quadrupole mass spectrometer (6460C, Agilent). Metabolites were separated using a gradient elution with mobile phases, 20 mM ammonium formate in H2O (phase A) and acetonitrile: water (90:10) at a final concentration of 20 mM ammonium formate (phase B), pH 3.0 for both mobile phases. A constant flow of 0.25 mL/min was used with a gradient of 100-90% B over 2 minutes and then 50% B over the next 6 minutes followed by returning to 100% B over 30 seconds before column re-equilibration for 6.5 minutes. The electrospray ionization source was operated with capillary voltage set to 3.5 kV, sheath gas temperature, 400 °C, sheath gas flow, 12 L/min, drying gas temperature, 350 °C, drying gas flow rate, 10 L/min, and nebulizer pressure, 60 psi. Sulfur metabolites were analyzed in positive mode by multiple reaction monitoring (MRM) of the transition from precursor to product ions at associated optimized collision energies and fragmentor voltages. Cell accelerator voltage was 4 V and dwell time was 5 ms. Two transitions were used for each compound, one being used for quantification and the second one for validation of identification. Data were analyzed using MassHunter Quantitative Analysis software. Absolute abundances were calculated by using external standard curves with normalized response factors using internal standards. Final concentrations were normalized to protein levels.

### Western blotting

Cells (500,000) were scrapped and collected, or tissues (∼20mg) were homogenized in 1x RIPA Lysis and Extraction Buffer (Thermo Scientific cat. no. 89900) supplemented with HALT protease inhibitor cocktail (Thermo Fisher Scientific, cat. no. 78430). Protein concentrations were determined by BCA assay (Thermo Fisher Scientific). Equal amounts of protein were loaded onto a 4-20% SDS-PAGE gel and transferred to a nitrocellulose membrane. The membrane was blocked in 5% bovine serum albumin in tris-buffered saline (TBS) for 2 hours at room temperature and incubated with primary antibodies at 4 °C overnight. Anti-Sds antibody (rabbit polyclonal, Genetex, cat. no. GTX47143, lot 82203710, 1:1,000) or Vinculin (mouse monoclonal, Abcam, cat. no. ab18058, lot GR287850-1, 1:1,000). The immunoblots were washed four times for 5 minutes with TBS-Tween (0.05%) incubated with polyclonal anti-rabbit IgG Alexa Fluor Plus 800 (Thermo Scientific, cat. no. A327325, lot XF349345, 1:10,000) and anti-mouse IgG Alexa Fluor 680 (Thermo Scientific, cat. no. A21058, lot 2478007, 1:10,000) secondary antibodies for 45 minutes at room temperature. Membranes were washed again in TBS-tween as described above and with TBS for 5 minutes. Blots were imaged with a LICORbio Odyssey CLx Imaging System equipped with Image Studio software.

### Serine palmitoyltransferase activity assay using microsomes

The microsome isolation protocol was adapted from a previously described report^63^. Two confluent 15-centimeter plates of HCT116 cells were scraped and pelleted in PBS. PBS was aspirated and the pellet was resuspended in 750 μL buffer A (10 mM HEPES-KOH, 1.5 mM MgCl2, 10 mM KCl, 1 mM EDTA, 1 mM EGTA, and 250 mM sucrose, pH 7.6) with protease inhibitor (Cat# 04693124001,Roche). Cells were pelleted and resuspended again in buffer A and left on ice for 30 minutes with manual inversions every 5 minutes. Cell membranes were ruptured by passing the suspension through a 22G needle and the nuclei were separated by centrifugation at 890 RCF for 5 minutes at 4 °C. The supernatant was collected and microsomes were pelleted at 20,000 RCF for 20 minutes at 4 °C. The microsome pellet was resuspended in a 50 mM HEPES (pH 8.1) and 2.5 mM EDTA buffer and stored at -80 °C. Microsomal protein content was quantified by BCA protein assay.

The SPT activity assay was adapted from previously described protocols^64^. Individual SPT reactions were performed at 37 °C with 100 μg of microsomal protein in a buffer containing 50 mM HEPES (pH 8.1), 50 μM PLP, and 2.5 mM EDTA. Reactions were initiated by the addition of 100 μM palmitoyl-CoA (Cat# P9716, Sigma Aldrich) and varying concentrations of serine from 0 to 40 mM. At 10 minutes, 100 μL of NH4OH and 100 μL of 1 M NaBH4 were added to catalyze the conversion of 3-ketodihydrosphingosine to sphinganine. At 20 minutes, the reaction was quenched by addition of 500 μL of methanol-chloroform (2:1, v/v) containing 20 pmol sphinganine-d7 internal standard followed by an additional 250 μL of chloroform and 500 ul of 0.5 M NH_4_OH. The samples were vortexed and centrifuged for 5 minutes and the upper aqueous layer was removed. The lower organic phase was washed with 500 μL of water, vortexed and centrifuged again, and then isolated and dried down. Total sphinganine d18:0 was quantified by an Agilent 6460C QQQ LC-MS equipped with a C18 column (Hypersil GOLD aQ C18 100 x 2.1 mm, 1.9 mm particle size, Thermo Fisher Scientific) as previously described^61^. Sphinganine was normalized to internal standard and reaction rates were calculated as nanomoles sphinganine/mg protein/minute. K_m_ and V_max_ were determined by performing a non-linear regression with the built- in Michelis-Menten function in GraphPad Prism 10.5.0.

### Statistical analysis

Data are presented as mean ± standard error of mean (SEM) of at least three biological replicates as indicated in figure legends. Statistical analysis was performed with GraphPad Prism 10.4.2 using Mann-Whitney non-parametric t-tests to compare two groups, one-way ANOVA with Kruskal-Wallis test to compare several groups and two-way ANOVA with Bonferroni’s multiple comparisons test for multiple comparisons within two groups. For all tests, p< 0.05 was considered significant with *p < 0.05, **p < 0.01, *** p< 0.001, or **** p<0.0001. Pearson’s correlation analysis was performed to assess linear associations between variables, and statistical significance was determined using two-tailed p-values.

## Acknowledgements

We thank all members of the Metallo Lab for helpful discussions and feedback. We acknowledge support from NIH grant R01CA234245, the Salk NCI Cancer Center CCSG P30CA014195, and the Lowy Medical Research Institute (all to C.M.M), the Salk Glenn Center, Paul F. Glenn Center for Biology and Aging Research (E.J.), R01DK111866, R56DK088837, DK099205, AA028550, DK101737, AA011999, DK120515, AA029019, DK091183, P42ES010337, R44DK115242 (all to T.K. and D.A.B.), R01CA285997 (D.A.B.), Stem Cell Fitness and Space Medicine Center at Sanford Stem Cell Institute (UCSD) (T.K.), and the NOMIS Foundation and the National Institutes of Health (R01-AI107027, R01-AI151123, R21-AI178938, R21-AI188938, and S10-OD034268) (all to Y.Z.). This work was supported by the GT3 Core Facility of the Salk Institute with funding from NIH-NCI CCSG: P30 CA014195.

## Author contributions

E.J. and C.M.M. designed the study. E.J., E.L.A., P.T., C.A., Y.L., M.L.R. performed animal experiments. E.L.A. conducted human cell line experiments. E.L.A. and P.T. performed enzymatic assays. E.J., E.L.A., C.A., A.L., and M.K.H., generated and analyzed targeted metabolomic data. J.R. generated plasmids. R.J. and M.L.G. performed human STT. H.Y.K. and T.K. provided donor samples. P.M., D.A.B., T.K., Y.Z., M.L.G., and C.M.M. guided experimental design and analysis. E.J. and C.M.M. wrote the manuscript with input from all authors.

---

**Extended Data Fig. 1.**
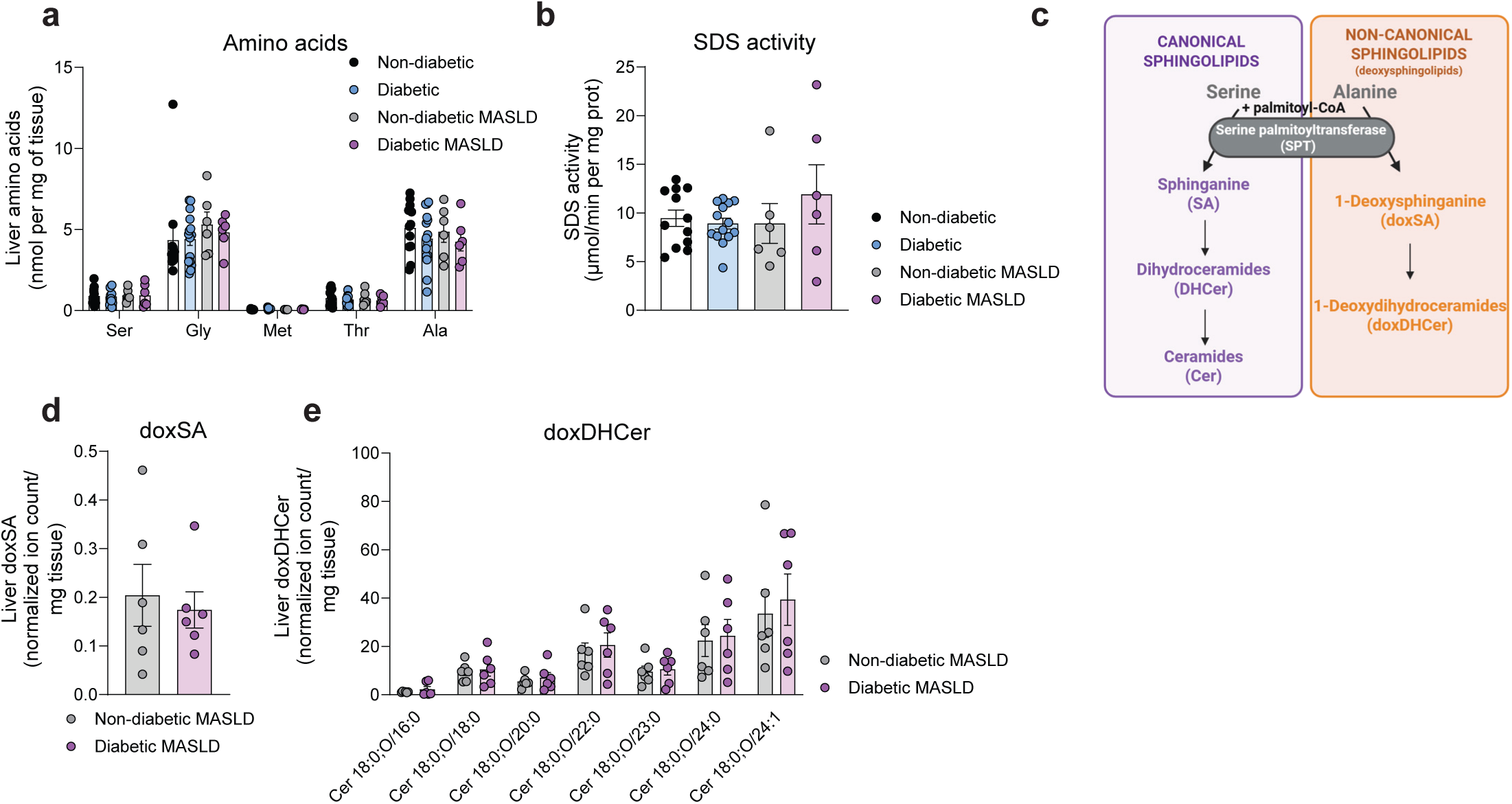
Serine dehydratase is expressed and active in human liver tissues. **a-b,** Liver amino acid concentration (**a**) and SDS activity (**b**) in non-diabetic (n = 12), diabetic (n = 15), non-diabetic MASLD (n = 6) and diabetic MASLD liver tissues (n = 6). **c,** Schematic of the sphingolipid biosynthesis pathway with canonical and non-canonical sphingolipids (created with BioRender). **d,e,** doxSA 18:0 (**d**) and doxDHCer 18:0 (**e**) relative abundance in non-diabetic (n = 6) and diabetic MASLD liver samples (n = 6). Data are presented as mean ± standard error of mean (SEM). Statistical analyses were performed using two-way ANOVA with Bonferroni’s multiple comparisons test (**a,e),** one-way ANOVA with Kruskal-Wallis test (**b)** and Mann-Whitney non-parametric t-test (**d**).

**Extended Data Fig. 2.**
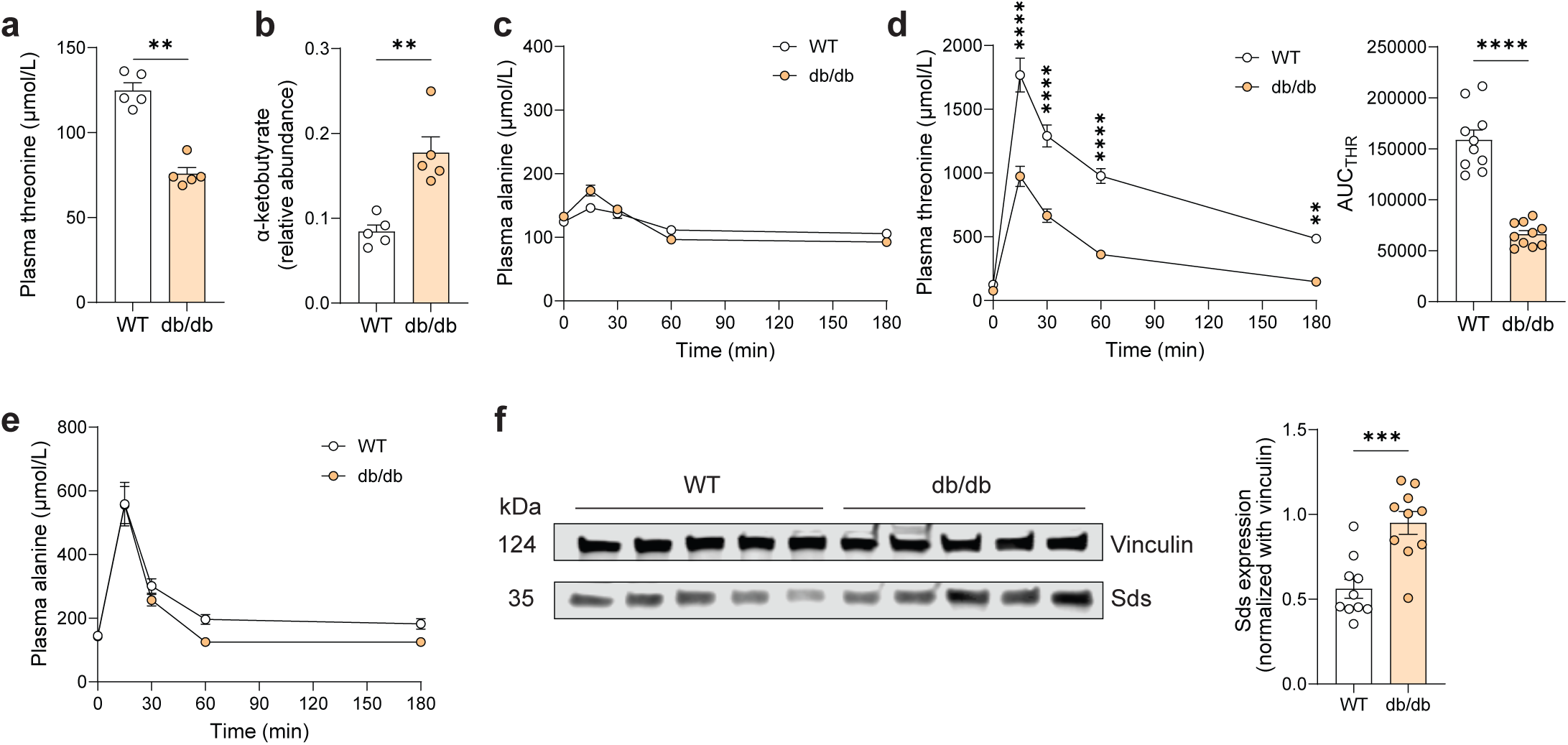
Threonine catabolism is increased in diabetic mice. **a,b,** Plasma threonine concentration (**a**) and alpha-ketobutyrate relative abundance (**b**) in control and diabetic mice (n = 5 per group). **c,** Plasma alanine concentration upon serine tolerance test (n = 12 per group). **d,** Plasma threonine concentration upon threonine tolerance test in WT and db/db mice (n = 10 per group). **e,** Plasma alanine concentration upon alanine tolerance test in WT and db/db mice (n = 6 per group). **f,** Representative Sds expression by western blot in WT and db/db mouse livers (left) (n = 5 per group) and quantification (right) (n = 10 per group). Data are presented as mean ± standard error of mean (SEM). Statistical analyses were performed using Mann-Whitney non-parametric t-test (**a**,**b**,**d** right, **f** right) and two-way ANOVA with Bonferroni’s multiple comparisons test (**c,d** left**,e**).

**Extended Data Fig. 3.**
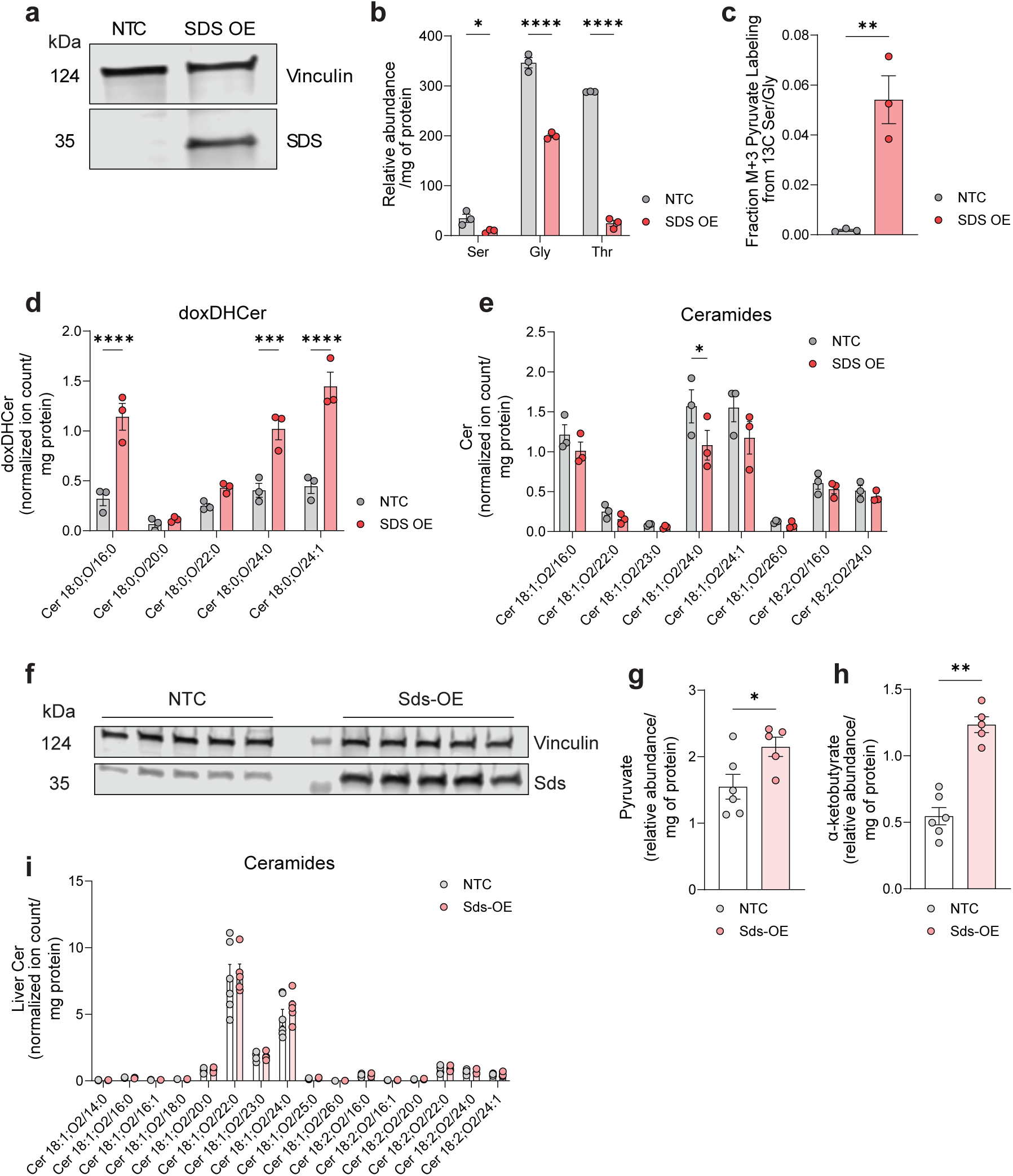
Sds overexpression promotes serine and threonine catabolism and alters sphingolipid biosynthesis. **a,** Western blot of human SDS in HCT116 cells treated with NTC or human SDS OE lentivirus following puromycin selection (n = 2). **b,** Relative abundance of serine, glycine, and threonine in HCT116 NTC and SDS OE cells for 72 hours (n = 3). **c,** Fraction of pyruvate with M+3 carbon labeling in HCT116 NTC and SDS OE cells cultured with [^13^C_3_]serine and [^13^C_2_]glycine for 6 hours (n = 3). **d,e,** Intracellular abundances of doxDHCer 18:0 (**d**) and ceramides 18:1 and 18:2 (**e**) in HCT116 NTC and SDS OE cells cultured for 72 hours (n = 3). **f,** Sds expression by western blot 4 weeks after AAV injection (n = 5 per group). **g-i,** Pyruvate (**g**), alpha-ketobutyrate (**h**) and ceramides 18:1 and 18:2 (**i**) relative abundance in NTC (n = 6) and Sds-OE mouse liver (n = 5), 4 weeks after AAV injection. Data are presented as mean ± standard error of mean (SEM). Statistical analyses were performed using two-way ANOVA with Bonferroni’s multiple comparisons test (**b,d,e** and, **i)** and Mann-Whitney non-parametric t-test (**c**,**g**,**h**).

**Extended Data Fig. 4.**
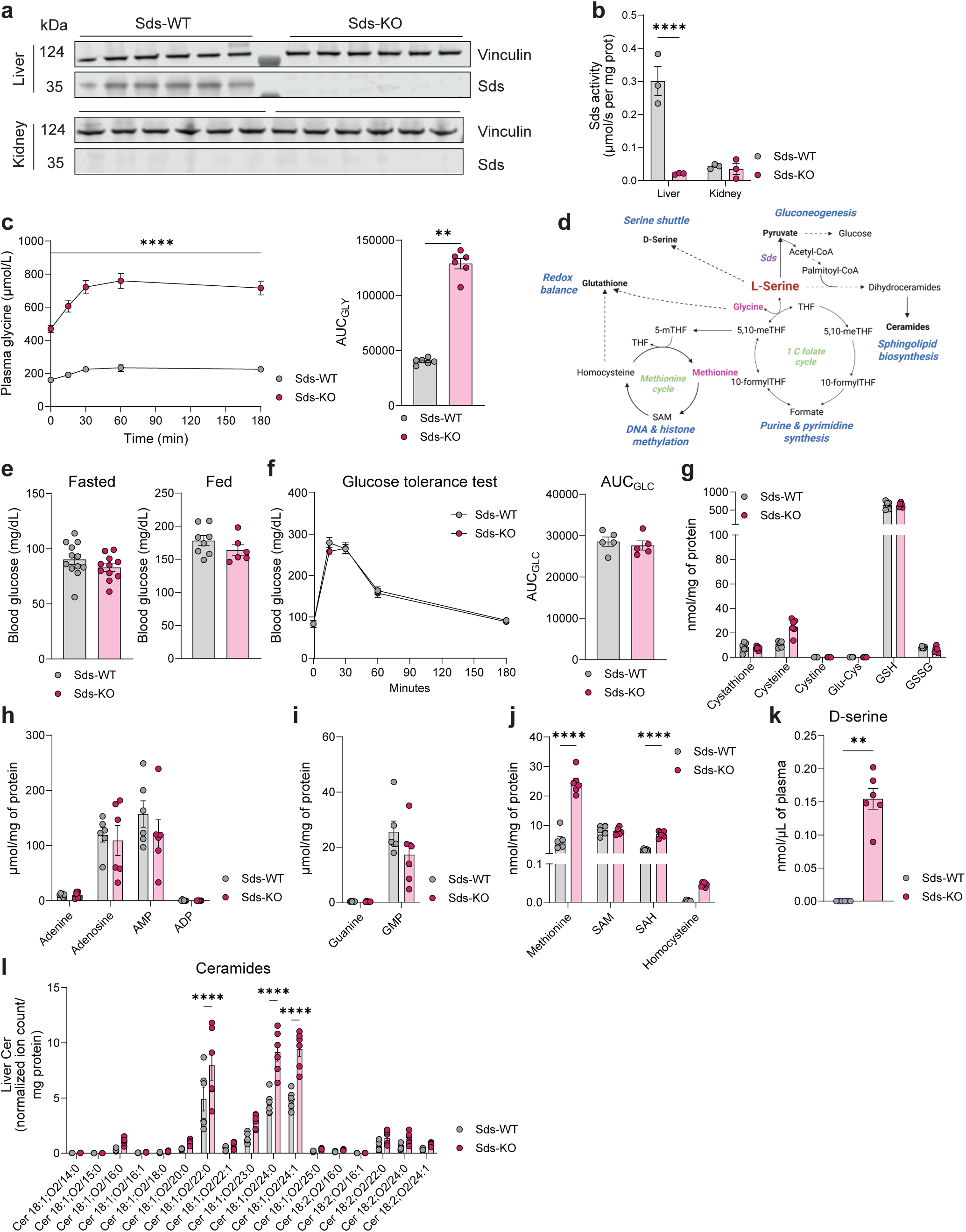
Sds deletion predominantly affects sphingolipid biosynthesis. **a,b,** Serine dehydratase expression by western blot (**a**) and activity (**b**) in 10–12-week-old Sds-WT and KO mouse liver and kidney (a, n = 6 per group and b, n = 3 per group). **c,** Plasma glycine upon serine tolerance test performed on 8-week-old Sds-WT and KO mice (n = 6 per group). **d,** Metabolic pathways downstream L-serine (created with BioRender). **e,** Fasted and fed blood glucose concentration in male and female Sds-WT and KO mice. **f,** Glucose tolerance test performed on male and female 10–12-week-old Sds-WT and KO mice (n = 5 per group). **g-j,** Glutathione pathway metabolites (**g**), purines (**h,i**) and methionine cycle metabolite (**j**) concentrations in Sds-WT and KO mouse livers (n = 6 per group). **k,** Plasma D-serine concentration in Sds-WT and KO mice (n = 6 per group). **l,** Liver ceramides 18:1 and 18:2 abundance in 10–12-week-old mice (n = 6 per group). Data are presented as mean ± standard error of mean (SEM). Statistical analyses were performed using two-way ANOVA with Bonferroni’s multiple comparisons test (**b,c** left, **f** left and, **g-j, l**) and Mann-Whitney non-parametric t-test (**c** right, **e**, **f** right and, **k**).

**Extended Data Fig. 5.**
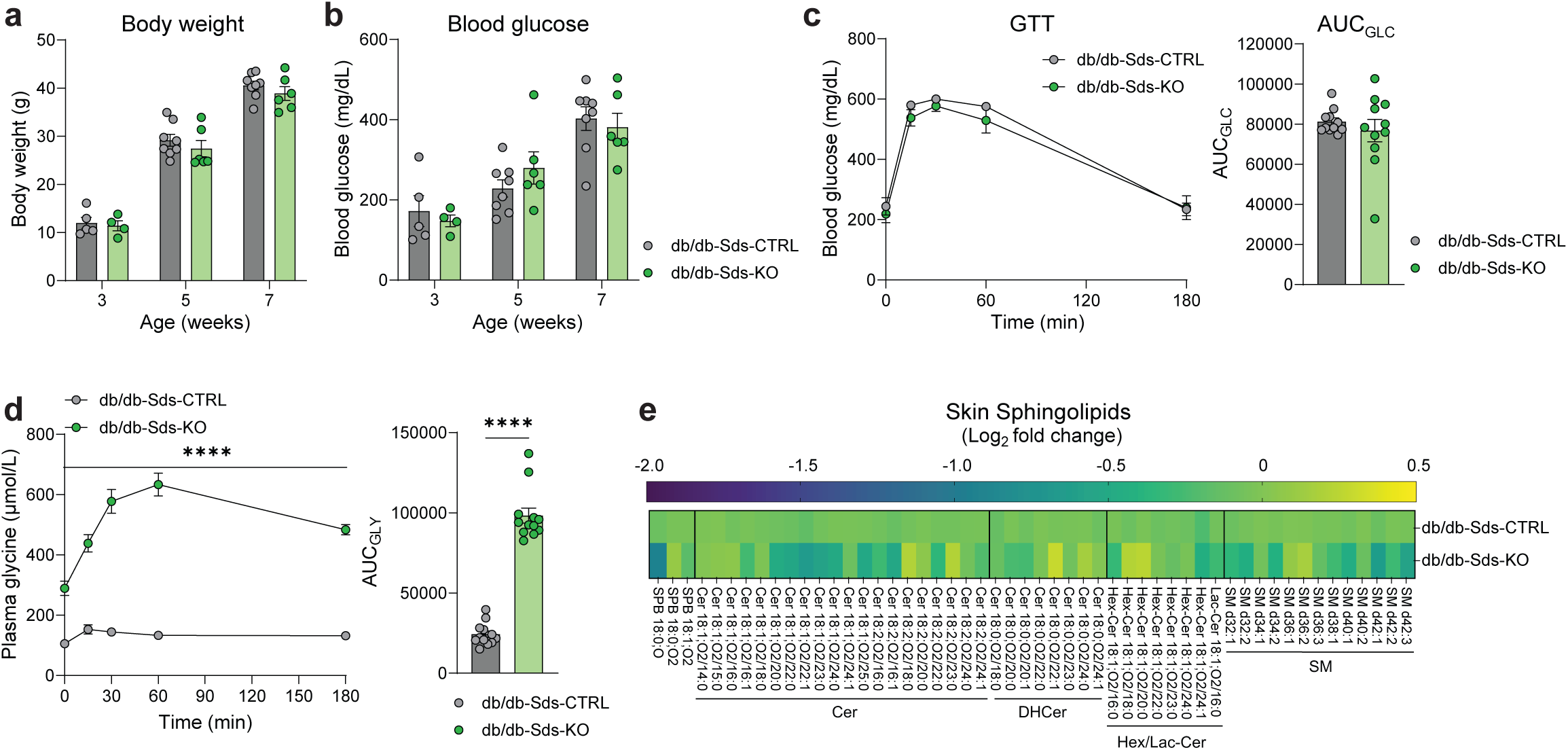
Obesity and diabetes are not affected by Sds deletion in db/db mice. **a,b,** Body weight (**a**) and blood glucose concentration (**b**) in 3-, 5- and 7-week-old female and male db/db-Sds-CTRL (n = 8) and KO mice (n = 6). **c,** Blood glucose upon glucose tolerance test on 10- to 14-week old female and male db/db-Sds-CTRL and KO mice (n = 11 per group). **d,** Plasma glycine concentration upon serine tolerance test performed on 10-14-week-old female and male mice (n = 12 per group). **e,** Skin sphingolipid profile in db/db-Sds-CTRL (n = 8) and KO mice (n = 6) - Cer: Ceramides, DHCer: Dihydroceramides, Hex/Lac-Cer: Hexosyl/Lactosyl-Ceramides, SM: Sphingomyelin. Data are presented as mean ± standard error of mean (SEM). Statistical analyses were performed using two-way ANOVA with Bonferroni’s multiple comparisons test (**a,b,c** left, **d** left and, **e**) and Mann-Whitney non-parametric t-test (**c** right and **d** right).

