## Supplementary Table 1 for "Targeting serine dehydratase supports amino acid homeostasis and skin repair"

**Supplementary Table 1: AAV and lentivirus plasmid sequences**

| **Organism** | **Gene** | **Plasmid info** | **Sequence** |
| --- | --- | --- | --- |
| *Homo sapiens* | GFP | pEGIP, puromycinR | MVSKGEEDNMAIIKEFMRFKVHMEGSVNGHEFEIEGEGEGRPYEGTQTAKLKVTKGGPLPFAWDILSPQFMYGSKAYVKHPADIPDYLKLSFPEGFKWERVMNFEDGGVVTVTQDSSLQDGEFIYKVKLRGTNFPSDGPVMQKKTMGWEASSERMYPEDGALKGEIKQRLKLKDGGHYDAEVKTTYKAKKPVQLPGAYNVNIKLDITSHNEDYTIVEQYERAEGRHSTGGMDELYK* |
| *Homo sapiens* | SDS | pEGIP, puromycinR | MMSGEPLHVKTPIRDSMALSKMAGTSVYLKMDSAQPSGSFKIRGIGHFCKRWAKQGCAHFVCSSAGNAGMAAAYAARQLGVPATIVVPSTTPALTIERLKNEGATVKVVGELLDEAFELAKALAKNNPGWVYIPPFDDPLIWEGHASIVKELKETLWEKPGAIALSVGGGGLLCGVVQGLQEVGWGDVPVIAMETFGAHSFHAATTAGKLVSLPKITSVAKALGVKTVGAQALKLFQEHPIFSEVISDQEAVAAIEKFVDDEKILVEPACGAALAAVYSHVIQKLQLEGNLRTPLPSLVVIVCGGSNISLAQLRALKEQLGMTNRLPK* |
| *Mus musculus* | mCherry | pZac | MVSKGEEDNMAIIKEFMRFKVHMEGSVNGHEFEIEGEGEGRPYEGTQTAKLKVTKGGPLPFAWDILSPQFMYGSKAYVKHPADIPDYLKLSFPEGFKWERVMNFEDGGVVTVTQDSSLQDGEFIYKVKLRGTNFPSDGPVMQKKTMGWEASSERMYPEDGALKGEIKQRLKLKDGGHYDAEVKTTYKAKKPVQLPGAYNVNIKLDITSHNEDYTIVEQYERAEGRHSTGGMDELYK* |
| *Mus musculus* | Sds | pZac | MAAQESLHVKTPLRDSMALSKLAGTSVFLKMDSSQPSGSFKIRGIGHLCKMKAKQGCRHFVCSSAGNAGMATAYAARRLGIPATIVVPNTTPALTIERLKNEGATVEVVGEMLDEAIQVAKALEKNNPGWVYISPFDDPLIWEGHTSLVKELKETLSAKPGAIVLSVGGGGLLCGVVQGLREVGWEDVPIIAMETFGAHSFHAAIKEGKLVTLPKITSVAKALGVNTVGAQTLKLFYEHPIFSEVISDQEAVSALEKFVDDEKILVEPACGAALAAVYSRVVCRLQDEGRLQTPLASLVVIVCGGSNISLAQLQALKVQLGLNGLPE* |
